# Strain typing and characterization of virulence genes in clinical *Staphylococcus aureus* isolates from Kenya

**DOI:** 10.1101/390955

**Authors:** Cecilia Kyany’a, Justin Nyasinga, Daniel Matano, Valerie Oundo, Simon Wacira, Willie Sang, Lillian Musila

## Abstract

*Staphylococcusaureus* strain typing is an important surveillance tool as particular strains have been associated with virulence and community and hospital acquired MRSA outbreaks globally. This study sought to determine the circulating strain types of *S.aureus* in Kenya and establish the virulence genes among the strains. Clinical *S.aureus* isolates from 3 hospitals in Kenya were sequenced on the Illumina Miseq and genomes assembled and annotated on PATRIC. Results demonstrated great diversity among the isolates with identification of 6 distinct CC (8,22,15,80,121,152), 8 ST types (8, 15, 22,80,121,152,241, 1633) and 8 spa types (t005, t037, t064, t084, t233, t2029, t272,t355). Novel STs (4705, 4707) and a novel spa type (t17826) were identified. The most prominent clonal complex was CC 152 comprised of only MSSA. A majority of MRSA isolates (3/4) typed to ST 241, CC8. One MRSA isolate typed to a novel ST 4705. All isolates were screened for a panel of 56 known virulence genes (19 adhesins, 9 hemolysins, 5 immune evasion proteins, 6 exo-enzymes and 19 toxins). 9 toxin genes were detected among the isolates with CC8 isolates having the highest numbers of toxin genes. An MSSA isolate (CC8) from a severe burn infection had the highest number of toxin genes (5). All MRSA isolates (CC8) had only 2 toxins, SEK and SEQ, whereas a majority of the MSSA isolates either had 0 or ≥2 toxins. SEK+SEQ and TSST-1+SEB+SEL toxin combinations were observed among patients whose disease resulted in hospitalization, an indicator of severe infections. This study confirms the highly heterogeneous *S.aureus* population in Kenya. MSSA appear to have the potential of accumulating more toxin genes than MRSA. This co-occurrence of major toxin genes, some associated with MRSA, highlights the potential risks of outbreaks of highly virulent MRSA infections which would pose treatment challenges.

## Introduction

*Staphylococcus aureus* is one of the leading causes of nosocomial infections but in recent years it has been increasingly associated with community acquired infections [1]. *S.aureus* causes a wide spectrum of diseases including bacteremia, pneumonia, urinary tract, skin and soft tissue infections (SSTI) [2,3]. Since the emergence of methicillin resistance (MRSA) in the 1940s epidemics initiated by successful MRSA clones have been observed [4,5]. For instance, USA 300, a highly virulent MRSA strain that emerged in the USA is currently associated with community outbreaks globally [6]. E-MRSA 15 which emerged in the UK has been linked to various hospital outbreaks [7]. Clonal success has been attributed to factors that enhance binding to host tissues and to the acquisition of virulence genes. For example, USA 300 has acquired the arginine catabolic mobile element, *sek* and *seq* virulence genes [8–10].

Strain typing is necessary for identification of emerging and outbreak associated strains. Multi locus sequence typing (MLST) and typing of the staphylococcal protein A (spa) gene have been widely used over the last 10 years to identify different strain types [11,12]. MLST examines differences in 7 housekeeping genes, assigns allele numbers and sequence types (ST) from the allelic profiles. These ST are then grouped into larger groups known as clonal complexes [13]. Spa typing is based on the number, sequence and type of repeats in the hypervariable region of protein A. For MRSA, additional typing of the staphylococcal cassette chromosome (SSC mec) which harbors the gene encoding methicillin resistance allows for additional discrimination between MRSA strains. Staphylococcal cassettes differ in gene arrangement and at least 11 types have been reported [14–16].

The *S.aureus* genome bears a plethora of virulence determinants [17,18] some of which are encoded in the core genome while others are borne on accessory genomes such as plasmids, conjugative transposons, plasmids and cassettes [19]. Virulence factors are grouped according to function into adhesins, immune evasion proteins, toxins and pore forming proteins. Cell wall adhesins mediate binding to host tissues and biofilm formation e.g. clumping factor A and polysaccharide intracellular adhesin. Pore forming proteins include leukocidins (LukE, LukD, and PVL) and hemolysins (gama, beta, alpha). *S.aureus* toxins have been associated with severe food poisoning (SEB, SEA) exfoliative skin conditions (ETA, ETB) and systemic shock (TSST-1) [20–24]. The distribution of virulence genes has been shown to differ between strain types [25].

Previous Kenyan studies have largely focused on susceptibility profiles [26,27] of *S.aureus* and strain typing with limited testing for virulence determinants which can influence infection severity. Virulence genes reported in both *S.aureus* carriage and infection studies in Kenya include Panton Valentine leukocidin (*PVL*), Toxic shock syndrome (TSST1), exfoliative toxin A and enterotoxin A with a notably high prevalence of *PVL* reported [28,29]. These studies were limited to 4 healthcare institutions in close geographic proximity therefore there is limited information on the diversity and distribution of the *S.aureus* population across Kenya. This study sought to fill this gap by typing isolates from a wider geographical area and identifying the relationships between Kenyan strains and known global strains using phylogeny. In addition, isolates were screened for a panel of known virulence genes and attempts made to correlate their presence with disease severity. By broadening our understanding of the *S.aureus* population in Kenya, this study provides baseline data for tracking emerging hypervirulent or outbreak associated strains of *S.aureus*.

## Materials and methods

### Ethics

This study was approved by the Walter Reed Army Institute of Research (#2089) and Kenya Medical Research Institute (#2767) IRBs.

### Bacterial isolates identification

Clinical *S.aureus* isolates from patients enrolled in an ongoing surveillance study (WRAIR#2089, KEMRI#2767) in public hospitals in three counties (Kisumu, Kericho and Nairobi) in Kenya were analyzed for this study. *S.aureus* isolates were identified based on beta hemolysis on sheep blood agar plates, gram positive, clustered cocci by gram stain, catalase and coagulase positive phenotypes. Isolate identity was confirmed using the MALDI-TOF biotyper (Bruker Daltonics, Millerica, MA, USA) and using the GP card on the Vitek 2 platform (bioMérieux, Hazelwood, MO, USA). 19 isolates (4 MRSA, 15 MSSA) obtained between April 2015 and December 2015 were studied. A majority of the *S.aureus* isolates were from skin and soft tissue infections (18/19). 84.2% were community acquired infections and 15.8% (3/19) were hospital acquired infections as per the CDC classification (29). Outpatient infections were considered mild infections (10/19, 52.6%) whereas inpatient infections (9/19, 47.3%) were considered severe infections, Table 1.

**Table 1:** Clinical characteristics of the Kenyan *S.aureus* isolates in this study.

| ISOLATE # | INFECTION TYPE<sup>d</sup> | MRSA/MSSA<sup>b</sup> | CA/HA<sup>c</sup> | IP/OP<sup>a</sup> | SEVERITY |
| --- | --- | --- | --- | --- | --- |
| SAKEN01 | UTI | MRSA | CA | OP | Mild |
| SAKEN05 | SSTI | MRSA | HA | IP | Severe |
| SAKEN06 | SSTI | MRSA | CA | IP | Severe |
| SAKEN13 | SSTI | MSSA | CA | OP | Mild |
| SAKEN14 | SSTI | MSSA | HA | IP | Severe |
| SAKEN15 | SSTI | MSSA | HA | IP | Severe |
| SAKEN16 | SSTI | MSSA | CA | OP | Mild |
| SAKEN17 | SSTI | MSSA | CA | OP | Mild |
| SAKEN18 | SSTI | MSSA | CA | IP | Severe |
| SAKEN19 | SSTI | MSSA | CA | IP | Severe |
| SAKEN20 | SSTI | MSSA | CA | IP | Severe |
| SAKEN21 | SSTI | MRSA | CA | IP | Severe |
| SAKEN22 | SSTI | MSSA | CA | OP | Mild |
| SAKEN23 | SSTI | MSSA | CA | IP | Severe |
| SAKEN24 | SSTI | MSSA | CA | OP | Mild |
| SAKEN25 | SSTI | MSSA | CA | OP | Mild |
| SAKEN26 | SSTI | MSSA | CA | IP | Severe |
| SAKEN27 | SSTI | MSSA | CA | OP | Mild |
| SAKEN29 | SSTI | MSSA | CA | OP | Mild |

Table legend: <sup>a</sup>IP-Inpatient, OP-Outpatient. Inpatient infections are considered severe infections in this100 study. <sup>b</sup>Methicillin Resistant *Staphylococcus aureus* (MRSA), Methicillin Susceptible *Staphylococcus aureus*.101 <sup>c</sup>CA – Community acquired infection, HA - Hospital associated infection. <sup>d</sup>SSTI – Skin and soft tissue
infection, UTI – Urinary tract infection

### Isolate typing

In-vitro spa typing was performed using conventional PCR [30] and Sanger sequencing. Contigs were assembled on CLC bio Main-Workbench (CLC bio, Aarhus, Denmark) and spa types analyzed using Ridom StaphType (Ridom GmbH, Münster, Germany). In-silico spa typing was done by analyzing assembled genomes on the online analysis pipeline https://cge.cbs.dtu.dk/services/spatyper/. Spa types obtained by both methods were compared. Sequences of isolates with novel spa repeats were submitted to the Ridom Spa Server for assignment of spa type.

MLST sequence type (ST) was determined in-vitro using published primers [11]. Gene sequences for each of the 7 loci were queried against the *S.aureus* database, allele numbers obtained and allelic profiles analyzed on http://www.mlst.net/ [13] to assign ST. Isolates were grouped into clonal complexes using the BURST clustering algorithm available on http://eburst.mlst.net/, allowing a minimum of 6 identical loci for group definition. Sequences of novel ST were submitted to https://pubmlst.org/saureus/ for ST assignment.

### Staphylococcal cassette chromosome typing

Staphylococcal cassette types for the MRSA isolates were determined using previously published primers [14]. PCR products were visualized on agarose gels and SSC mec types determined based on different amplicon size.

### Whole genome sequencing and sequence analysis

Genomic DNA was extracted from freshly cultured *S.aureus* isolates using ZR Fungal/Bacterial DNA MiniPrep Kit (Zymo research, California, United States). DNA concentrations were determined using the Qubit (Thermo Fisher Scientific, Massachusetts, United States) and 1ng of DNA used for library preparation with the Nextera XT kit (Illumina) as per manufacturer’s instructions to generate 300bp paired end libraries. Libraries were sequenced on an Illumina MiSeq platform.

Raw reads were uploaded onto the Pathosystems Resource Integration Center (PATRIC 3.5) https://www.PATRICbrc.org/ [31]. Genome assembly was carried out on PATRIC using the assembly pipeline ‘miseq’ as the assembly strategy which runs both Velvet [32] and SPAdes [33] algorithms and uses ARAST, an in-house scoring script. Annotation was performed using the RASTk pipeline [34] on PATRIC with ‘Bacteria’ as the domain and ‘Staphylococcus aureus’ as the taxonomy ID. Genome assemblies were uploaded onto NCBI under BioProject ID PRJNA481322. A summary of the genome characteristics can be found in Table 2.

**Table 2:**
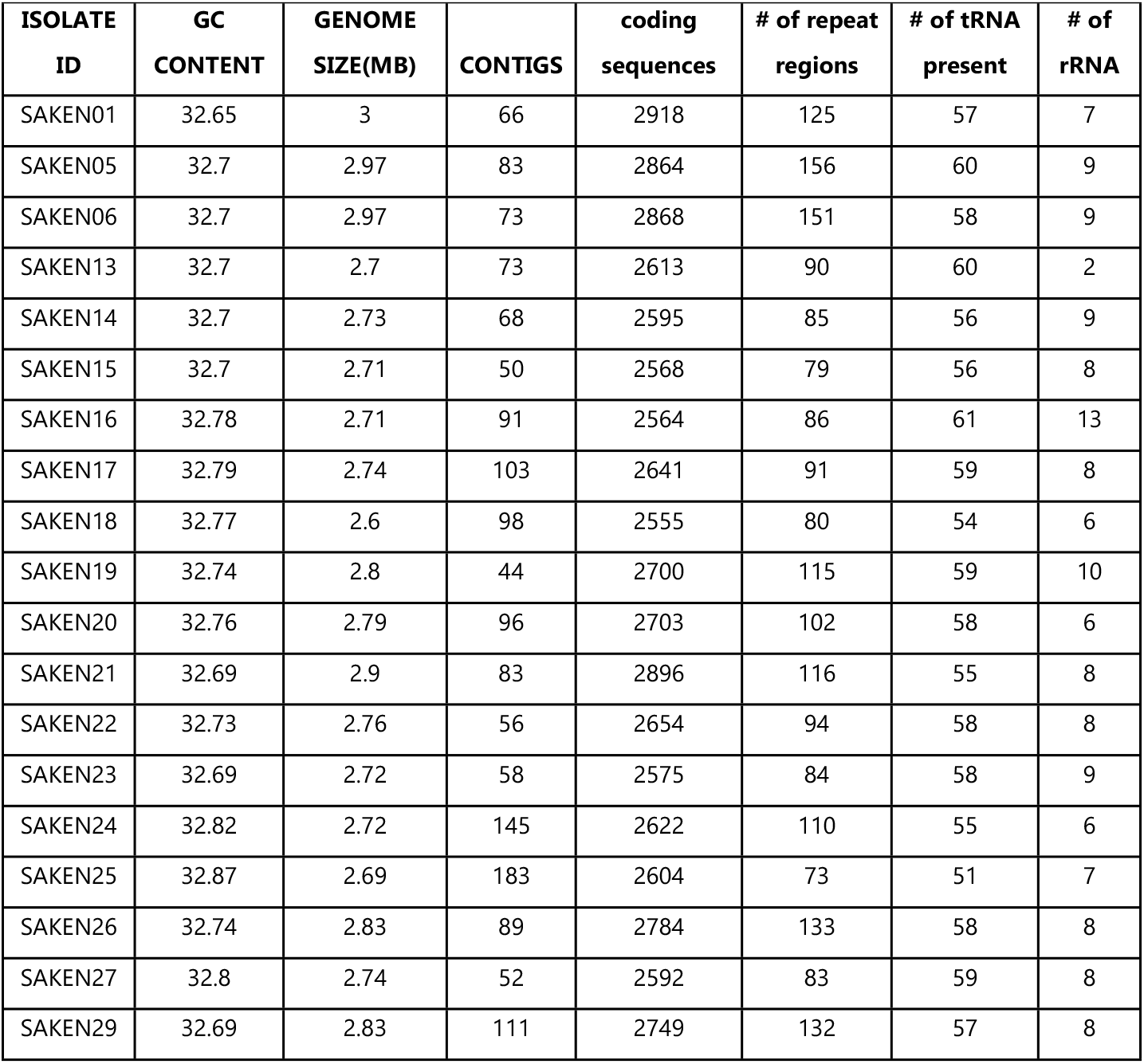
Genome characteristics of *S.aureus* isolates analyzed in this study.

### Bioinformatic analysis

In-silico MLST was determined by analyzing the assembly files on both the PATRIC genome characterization pipeline and on MLST 1.8 https://cge.cbs.dtu.dk/services/MLST/ [35] at the Centre for Genomic Epidemiology (CGE). Novel ST types were submitted to https://pubmlst.org/saureus/ for typing [36].

To establish the relationships between the isolates of this study, MLST sequences for the query genomes were concatenated on FASconCAT [37] and maximum likelihood phylogeny inferred using PhyML 3.1 [38] utilizing the TMP model with gamma variation.

To infer relationships between genomes of this study and known global strains, phylogenetic analysis was performed on the 19 Kenyan *S.aureus* whole genomes and 26 global strains. Reference strains were selected to include at least one whole genome for all the sequence types identified and common global and regional strains. The reference strains used in the phylogenetic analysis are under supplementary information appendix S1, Table 1. To generate a whole genome sequence phylogeny on PATRIC, genome assemblies were analyzed on the phylogenetic tree building service. The tree building service filtered genome protein files and used BLAST to determine the best bi-directional hit, clustered protein files into homolog sets and filtered out homolog sets representing <80% of the genomes and eventually protein sets meeting the threshold were trimmed and aligned with MUSCLE [39]. Tree alignments were concatenated and a main tree built using RAxML [40]. Progressive tree refinement was employed for analysis of poorly refined sub - trees [31,41]. Phylogenetic trees generated from both the MLST loci and WGS (Figure 1-2) were compared.

**Fig 1:**
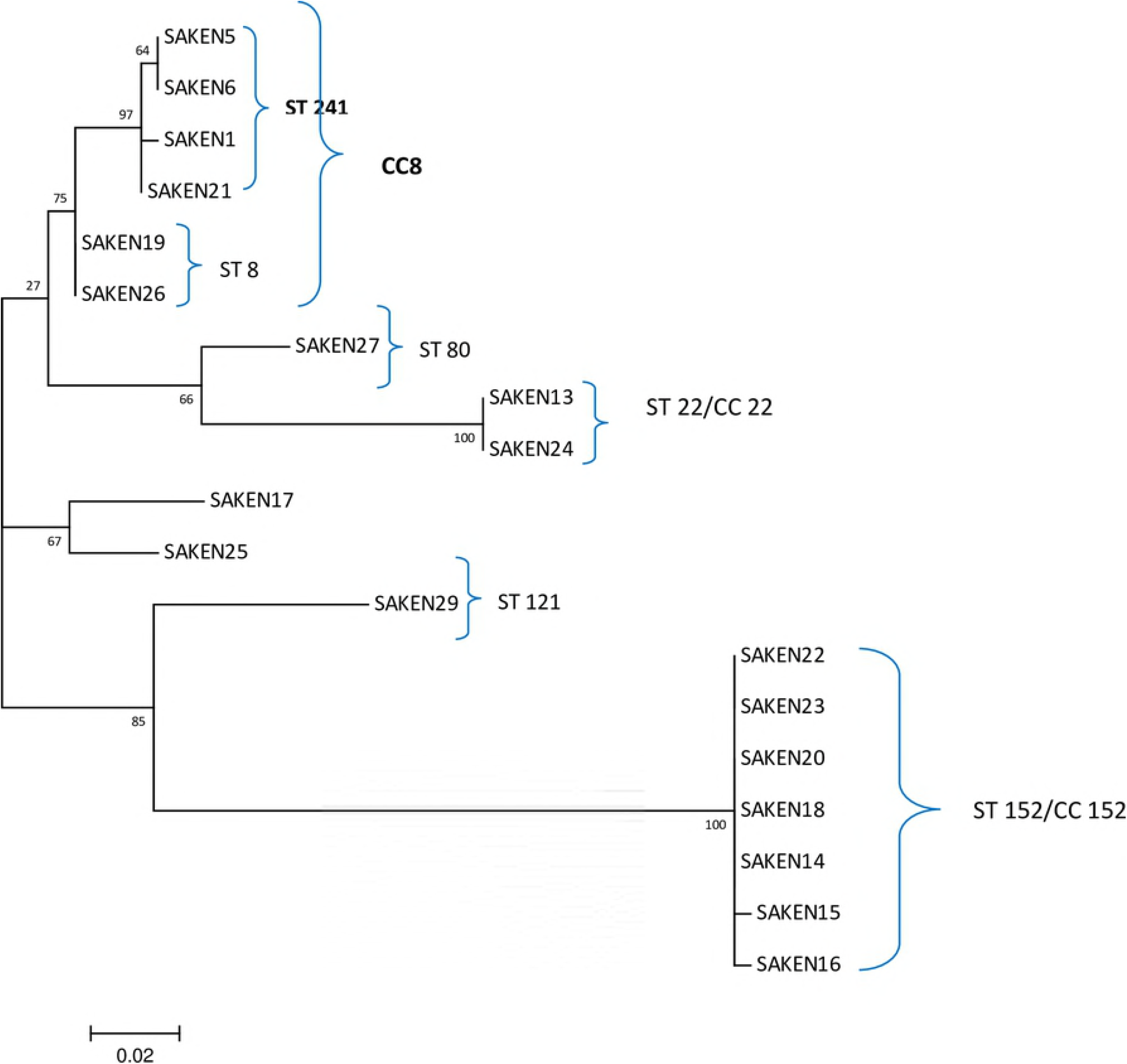
Dendogram of an MLST phylogeny of Kenyan *S.aureus* isolates. Maximum likelihood was phylogeny inferred using a custom model and 100 boot strap replicates. ST and CC associated with MRSA isolates of this study are depicted in bold font.

**Fig 2:**
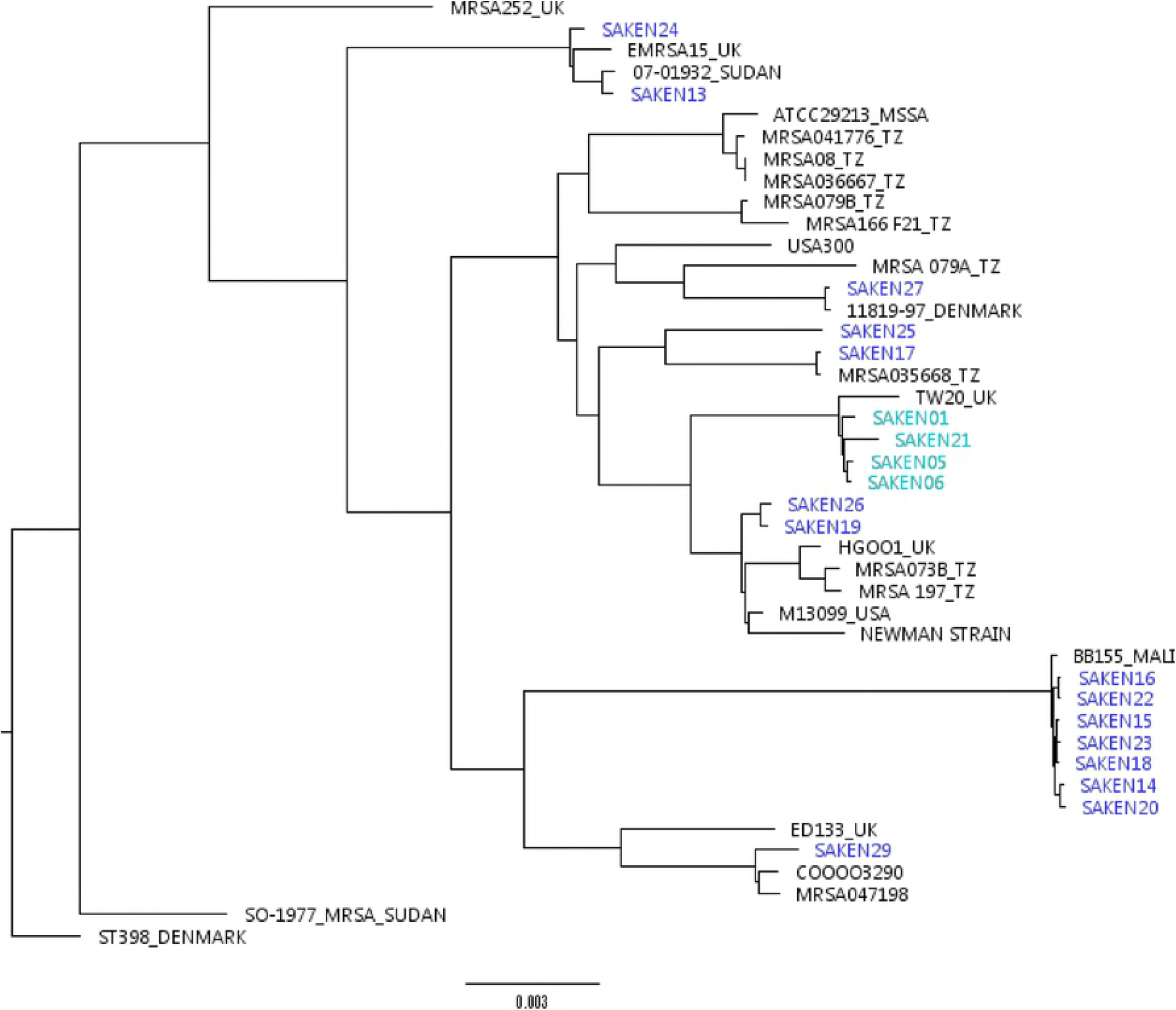
Dendogram showing whole genome phylogeny of Kenyan isolates. (SAKEN prefix and in blue) relative to known global strains. MRSA isolates are depicted in green. Reference isolates are in black with a country/regional prefix. Phylogeny inferred using maximum likelihood with progressive refinement on PATRIC.

To investigate the presence of known virulence genes, whole genomes of Kenyan isolates were screened for 56 virulence genes. Virulence genes were identified under the specialty genes on PATRIC with ‘Virulence’ as the search term. Targeted virulence genes and pathogenicity islands were identified across all isolates on PATRIC. To verify the virulence genes identified, contigs were analyzed on VirulenceFinder 1.5 https://cge.cbs.dtu.dk/services/VirulenceFinder/ [42]. The virulence genes identified were then grouped according to function.

## Results and Discussion

### Spa typing

Study isolates typed into 8 distinct spa types (t005, t037, t064, t084, t233, t2029, t272, t355). t355 was the dominant spa type (7/19; 36.8%). A single novel spa type, assigned t17826, was reported. MRSA isolates typed to t007 (1) and t037 (3) while MSSA isolates showed great spa diversity with t005, t17826, t272, t13194 being represented by singletons. Discrepancies between in-vitro and in-silico spa typing were observed for SAKEN 01, (t007 vs t2029) and SAKEN 13, (t084 vs t233) (Table 2).

### MLST typing

Multi locus sequence typing results were consistent on both CGE and PATRIC analysis platforms. There was great diversity among the isolates with the identification of 6 distinct CC (8,22,15,80,121,152) and 8 ST types (8, 15, 22,80,121,152,241, 1633) (Table 3). 2 novel STs assigned ST 4705 (CC 8, MRSA) and ST 4707 (CC 5, MSSA) by PubMLST [36] were reported. A majority of the isolates belonged to CC 152 (7/19) and CC 8 (5/19). ST 152 was the most dominant 6/19 (31%) with all isolates being MSSA geographically distributed across the 3 counties. 4 of the STs were represented by singletons.

**Table 3:** Table showing STs, CCs of Kenyan *S.aureus* isolates.

| Isolate | Clonal complex | MLST type (ST) | SPA type (in-vitro) | SPA type (in- silico) | MRSA/MSSA | REGION <sup>a</sup> |
| --- | --- | --- | --- | --- | --- | --- |
| SAKEN01 | 8 | 4705* | **t007** | **t2029** | **MRSA** | KDH |
| SAKEN05 | 8 | 241 | t037 | t037 | **MRSA** | KDH |
| SAKEN06 | 8 | 241 | t037 | t037 | **MRSA** | KDH |
| SAKEN21 | 8 | 241 | t037 | t037 | **MRSA** | KDH |
| SAKEN19 | 8 | 8 | t064 | t064 | MSSA | KDH |
| SAKEN26 | 8 | 8 | t064 | t064 | MSSA | KDH |
| SAKEN16 | 152 | 152 | t355 | t355 | MSSA | KCH |
| SAKEN18 | 152 | 152 | t355 | t355 | MSSA | KCH |
| SAKEN20 | 152 | 152 | t355 | t355 | MSSA | KDH |
| SAKEN14 | 152 | 152 | t355 | t355 | MSSA | NRB |
| SAKEN22 | 152 | 152 | t355 | t355 | MSSA | NRB |
| SAKEN23 | 152 | 152 | t355 | t355 | MSSA | NRB |
| SAKEN15 | 152 | 1633 | t355 | t355 | MSSA | NRB |
| SAKEN24 | 22 | 22 | t005 | t005 | MSSA | KCH |
| SAKEN13 | 22 | 22 | **t084** | **t233** | MSSA | NRB |
| SAKEN25 | 5 | 4707* | t17826* | unknown | MSSA | KDH |
| SAKEN17 | 15 | 15 | t084 | t084 | MSSA | KCH |
| SAKEN27 | 80 | 80 | t13194 | t13194 | MSSA | KCH |
| SAKEN29 | 121 | 121 | t272 | t272 | MSSA | KDH |

Table legend. \* denotes novel ST and spa types reported by this study. <sup>a</sup> Three regions in Kenya where the
isolates were obtained from: KCH- Kericho, KDH – Kisumu County, NRB- Nairobi County

All the MRSA in this study belonged to CC 8. A majority of MRSA (3/4) in this study typed to ST 241 and spa type t037. One isolate typed to the novel ST 4705 and spa type t007. This isolate only differed from the other isolates in the acetyl coenzyme A acetyltransferase (*yqil*) loci. In-vitro SSC mec typing identified 2 SSC mec types with a majority of the MRSA typing to SSC mec IVc (3/4). The remaining isolate with the novel ST 4705, typed to SSC mec II.

Each region showed a unique genetic fingerprint with CC 8 (ST 8 and ST 241) detected in Kisumu County only while CC 152 showed a wide geographic distribution. Kericho County showed greatest heterogeneity in CC/ST.

### Phylogenetic analysis

Maximum likelihood phylogenies reconstructed on PATRIC and on PhyML gave similar tree topologies with isolates clustering according to sequence types and/or clonal complexes Fig 1-2.

The phylogenetic tree inferred using MLST depicted a multifurcating tree with distribution of MRSA and MSSA isolates in distinct clusters. ST 121 was ancestral to ST 152 while ST 80 was ancestral to ST 22. CC 8 cluster encompassed all MRSA isolates (ST 241, ST 4705) and ST 8 MSSA isolates. The MRSA clade (ST 241) and MSSA (ST 8) clade are sister groups sharing a recent common ancestor. The MRSA isolate (SAKEN 01) which has a novel ST grouped within the MRSA clade.

Whole genome phylogeny with global strains showed Kenyan ST 152 isolates clustered with the asymptomatic Malian BB155 strain. MRSA in this study were closely related to TW20 strain 582 (ST 239) which is a successful HA MRSA lineage in the United Kingdom [7]. Kenyan ST8 MSSA isolates grouped closely with the highly virulent Newman strain [43]. ST/CC 22 was represented by 2 MSSA isolates which clustered with EMRSA-15 (MRSA-15), associated with widespread outbreak of HA-MRSA in Europe [44]. It was also observed that Kenyan strains in this study were closely related to MRSA and MSSA isolates from Tanzania and Sudan respectively. Tanzanian MRSA isolates clustered with the Kenyan ST 8 MSSA isolates and Kenyan ST 15 MSSA.

### Virulence genes screen

The 19 *S.aureus* genomes were screened in-silico for a panel of 56 known virulence genes (19 adhesins, 9 hemolysins, 5 immune evasion proteins, 6 exo-enzymes and 19 toxins). Of the 19 adhesins investigated, 10 were identified in all the isolates (ClfA, ClfB, FnbA, EbpS, IcaA, Eap/Map, SasA, SacC, SasF, and SasH) (S2 table). Since these adhesins are ubiquitously expressed as they are essential for successful host invasion by facilitating binding to extra cellular matrix proteins [45], they are not discussed further in this paper. EfB (Extracellular Fibrinogen – binding protein) was detected in 84% of the isolates. 18/19 of the isolates bore ≥2 of the Ser-Asp rich fibrinogen binding proteins whereas collagen binding protein (cna) was present in 63% of the isolates. It was observed that the presence of the adhesin genes was loosely correlated with the clonal complex. For example, isolates lacking Efb (3) belonged to CC 152 while all CC 152 isolates lacked SdrC, SasG cell wall adhesion protein and CC8 isolates lacked the SasK adhesion protein.

A majority of the hemolysins screened for (6/9) were identified in the isolates (gamma A, B, C, delta, alpha hemolysin and PVL). 63% (12/19) of the isolates had ≥5 of the hemolysins screened for with Leukocidin D and E being present in 94% and 89% of the isolates respectively. Hemolysins damage the host cell plasma membrane which is a critical process during *S.aureus* infection and disease progression [46]. Of note, all isolates typing to CC 152 lacked beta hemolysin whereas the bi-component protein Panton Valentine leukocidin was detected in all the Kenyan isolates.

Of the 19 toxins screened for 9 were detected; TSST-1, ETA/ETB, SEB, SEG, SEH, SEK, SEL, SEQ, and SPEG. Toxins were found to be randomly distributed among the CCs. TSST-1 was detected in 3/19 of the isolates obtained from various soft tissue infections (burns, abscesses and cuts). All the isolates with TSST-1 were MSSA of which 2/3 were from hospitalized patients an indicator of severe infection. Staphylococcal pathogenicity island 1 (SaP1) was detected in all isolates with TSST-1 (S2 Table 1). SEB was present in 2/19 of the isolates obtained from hospitalized patients presenting with a burn and an abscess. SEL, SEG enterotoxins and ETA/ETB exfoliative toxins were each detected in 2/19 of the isolates. Isolates with ETA and ETB exfoliative toxins were obtained from outpatient subjects with mild abscess and skin lesion infections. Staphylococcal pathogenicity island 2 (SaPln2) was detected in isolates with one or both exfoliative toxins.

Some toxin combinations were observed to always co-occur: SEK+SEQ, SEB+SEL, TSST-1+SEG, TSST-1+SEB+SEL, TSST-1+SEB+SEK+SEL. The SEK+SEQ combination was present in all the MRSA isolates in this study a majority (3/4) of which were from hospitalized subjects. TSST-1 detected in this study was from both hospitalized (2) and outpatient (1) subjects. In the hospitalized patients TSST-1 co-occurred with SEB and SEL while in an outpatient TSST-1 co-occurred with SEG suggesting that TSST-1+SEB+SEL combination could cause severe infections.

Comparison of the number of toxin genes between MRSA and MSSA isolates revealed that all Kenyan MRSA isolates only had 2 toxins whereas MSSA isolates had between 0-5 toxins. The isolate with the highest number of toxins (5) was an MSSA isolate obtained from a hospitalized burn patient. This MSSA isolate was in the same CC as the MRSA isolates suggesting the potential risk of emergence of CC 8 strains that are both highly virulent and difficult to treat. A majority (2/3) of the hospital acquired infections had no toxin genes detected in their genomes with the remaining isolate having 2 toxin genes.

## Discussion

This study typed *S.aureus* isolates from widespread geographical areas in Kenya and inferred phylogenetic relationships between the isolates and known global and regional strains. To understand the virulence potential of the Kenyan strains, virulence genes in the Kenyan strains were identified and the presence of these genes investigated in relation to CC and clinical presentation.

Strain typing revealed 8 STs and 8 spa types among the 19 Kenyan isolates confirming the great heterogeneity previously described among *S.aureus* both regionally and globally [28,29,47–49]. The isolates grouped into 6 distinct clonal complexes with CC 152 being most dominant and widely distributed across the three counties. CCs 8 and 22 known to harbor MRSA globally were also reported. All MRSA isolates in this study typed to CC 8 while MSSA isolates showed heterogeneous distribution across a number of CCs.

Globally, MRSA strains have been shown to belong to 3 major CC: CC 8, CC 5 and CC 22 [50]. For instance, EMRSA15/UK that is responsible for hospital acquired MRSA infections in the UK types to CC 22. In this study CC 22 was comprised of only MSSA isolates whereas CC 8 was composed of both MSSA and MRSA isolates. Previous Kenyan studies carried out in hospitals in Nairobi [47] and Thika [28] identified CC 5 as the predominant circulating MRSA clone within hospitals in Nairobi and its environs. MRSA isolates from hospitals in Nairobi were predominantly ST 241, t037 while ST 239, t037 prevailed in the hospital in Thika. In this study, all MRSA isolates typed to ST 241, t037; similar to that reported by Omuse et al [47] in 3 hospitals in Nairobi. All MRSA isolates in this study (CC 8, ST 241 and novel ST 4705) were obtained from Western Kenya which is situated 300km from Nairobi. This identification of ST 241 MRSA strains in both Western Kenya and Nairobi suggests a widespread geographical distribution of this MRSA strain in Kenya. Schaumburg et al [48] reported ST 241 MRSA clone to be widespread in Africa though with varying SSCmec types; Senegal (SSCmec III), Tunisia (SSCmec III), Niger (SSCmec III and V) Nigeria (SSCmec III and IV) and Algeria (SSCmec III) [51]. A majority of the MRSA isolates of this study bore SSCmev IV similar to that reported in Nigeria.

Spa typing showed a greater discriminatory power than ST with multiple spa types belonging to the same STs. Discrepancies between in-vitro and in-silico spa types (t007 vs t2029 and t084 vs t2330) was due to shorter repeat sequences in-silico. The shorter repeats observed in silico can be attributed to filtering out of low quality reads in the sequence analysis pipeline. Thus, we recommend the use of conventional in-vitro spa tying as a surveillance tool more so in regions with great *S.aureus* population diversity.

Inferred phylogeny of the Kenyan isolates showed distinctive clustering by CC. CC 8 cluster composed of two clades ST 8 (MSSA) and ST 241 (MRSA) with the two clades sharing a recent common ancestor. Studies have shown that MSSA isolates of CC 8 act as reservoirs for MRSA pending acquisition of the staphylococcal cassette [52,53].

Relationships between isolates of this study and known global strains using whole genome phylogeny revealed close clustering of MRSA strains in this study with the well-known TW20 strain 582, which is a successful HA MRSA clone which originated from London in the UK [7]. TW20 is a hospital associated outbreak MRSA strain known for its high transmissibility and multi-drug resistant properties due to a plethora of resistance genes carried on mobile elements [54]. TW20 and the MRSA strains in this study both type to CC 8. CC 8 MRSA strains have been linked to community acquired infections and is the predominant MRSA strain in this study. This clonal complex encompasses well known strains such as USA 300 which is a lineage linked to the acquisition of SSCmec IV, PVL and SEQ and SEK genes [10,55]. EMRSA-15 strain belongs to the same clade as Kenyan ST 22 MSSA isolates. EMRSA-15 is a well-known strain associated with the widespread outbreak of HA-MRSA in Europe [44] and MSSA isolates typing to ST 22 have been identified as the MSSA reservoir from which EMRSA-15 emerged [56]. ST 152, the dominant Kenyan strain is related to both carriage strains in Mali [57] and pathogenic strains in Europe [58–60].

Bacterial virulence factors are key for successful host colonization and infection. Adhesins facilitate successful binding to the host extra cellular matrix promoting subsequent biofilm formation. Worth noting is that ST 152 isolates in this study lacked some adhesins (SasG and SdrC) and hemolysin (beta) genes. SasG surface protein mediates successful bacterial adhesion to squamous epithelial cells of the nostrils and promotes biofilm formation. Expression of SasG has been shown to mask the effect of adhesins binding to ligands such as fibrinogen and fibronectin [61]. Both Malian BB155 and Kenyan ST 152 isolate genomes lack SasG which we speculate could promote successful host colonization and provide a fitness advantage. All CC8 isolates lacked SasK cell wall adhesion, the significance of this putative adhesin is yet to be determined [62]. Previous studies by McCarthy and Lindsay [25] noted similar observation of variation in surface adhesions between clonal complexes and lineages.

PVL, a bi-component leukocidin causing destruction of leukocytes and tissue necrosis, was widespread across all isolates in this study. PVL positive MRSA clones were identified in this study and in other Kenyan and African studies [27,29,63,64]. A Nigerian study pointed out the possibility of emergence of PVL positive MRSA clones as a result of the co-existence of MRSA clones and PVL positive MSSA [65]. This study reports extensive (100%) PVL presence in Kenyan MSSA and MRSA isolates supporting this hypothesis.

Staphylococcal toxin (SEB) is a super antigenic toxin associated with food poisoning, non-menstrual toxic shock syndrome, dermatitis and asthma [20]. Kenyan isolates bearing SEB typed to CC 8 and CC 152 consistent with previous observations where SEB was identified most often in CC8 isolates in New York [66]. A Taiwanese study identified SEB to be the cause of Staphylococcal scarlet fever [67]. In this study, SEB was detected in MSSA isolates obtained from a burn and from an abscess. In both cases these were inpatients indicative of severe infections.

Toxic shock syndrome toxin (TSST-1) was initially reported as the cause of menstrual toxic shock but over the years non-menstrual toxic shock syndrome has been reported [68–70]. TSST-1 is encoded by tstH gene borne on the staphylococcal pathogenicity island 1 [71]. SaP1 was identified in all the isolates positive for TSST-1. TSST-1 was observed to co-occur with up to 4 staphylococcal enterotoxins and most often with SEB and SEL. The presence of TSST-1 and SEB toxins in isolates obtained from severe infections resulting in hospital admissions suggests the severity of this toxin combination.

SEK and SEQ have been linked to a number of food poisoning cases [24,72]. Among the Kenyan isolates these enterotoxins were associated with SSTIs. SEQ+SEK combination was consistently observed in Kenyan MRSA isolates. SEQ and SEK toxin genes were reported to significantly cooccur in Chinese MRSA isolates [73]. These toxins co-occur on genomic islands and have been associated with the HA SSC mec II clone [21,66]. CC 8 isolates have been reported to consistently bear SEQ, SEL and SEK toxin genes [19]. This pattern of distribution was observed among isolates in this study.

Of the 4 MRSA isolates 3 were MRSA ST 241, mec IVc and the remaining isolate was the novel MRSA ST 4705, mec II. Worth noting was the difference between the SSC mec cassettes present in the isolates and that the MRSA ST 4705, mec II was the sole MRSA isolate associated with a community acquired UTI infection.

It was also observed that a majority of MRSA isolates from this study were community acquired and bore PVL, enterotoxin Q and K toxin genes. Studies by Voyich et al [74] have suggested the highly virulent nature of community acquired MRSA in comparison to hospital acquired strains a hypothesis that is not supported by this study which showed no differences in numbers and types of virulence genes between the hospital acquired and the community acquired strains.

## Conclusion

Despite the low number of isolates analyzed, this study provides a glimpse into the diversity and distribution of Kenyan MSSA and MRSA isolates and their relatedness to global strains. The study highlighted the potential impact of particular toxin combinations on clinical severity and provided evidence that co-occurrence of methicillin resistance and virulence genes could portend the emergence of highly virulent MRSA infections. There is demonstrated need for continued trend monitoring through surveillance which will continue as part of this ongoing surveillance program.

## Acknowledgements

We thank Dr John Waitumbi, Kimita Gathii and Luiser Ingasia from Kisumu basic Science Lab for sequencing support, Ann Wattam (PATRIC) and Frances Coll (Sanger Institute) for assistance in WGS analysis and phylogeny. We are grateful to all the study subjects in the participating healthcare facilities. This work has been published with the permission of the Director KEMRI.

Material has been reviewed by the Walter Reed Army Institute of Research. There is no objection to its publication. The opinions or assertions contained herein are the private views of the author, and are not to be construed as official, or as reflecting true views of the Department of the Army or the Department of Defense. The investigators have adhered to the policies for protection of human subjects as prescribed in AR 70–25.

## Author contributions

Conceptualization: LM

Data Curation: LM, CK

Formal analysis: CK, JN, LM

Funding Acquisition: LM

Investigation: CK, JN, DM, VO, SW

Methodology: LM

Supervision: VO, LM, WS

Validation: LM, CK

Visualization: LM, CK

Writing-Original draft preparation: CK, LM

Writing-Review and Editing: JN, CK, VO, SW, DM, WS, LM

## SUPPLEMENTARY INFORMATION

**S1 Table 1: List of reference genomes used in this study**

**S2 Table 2: Virulence gene profiles across *S. aureus* isolates**

## REFERENCES

1. Klein E, Smith DL, Laxminarayan R (2007) Hospitalizations and Deaths Caused by Methicillin-Resistant Staphylococcus aureus, United States, 1999–2005. Emerg Infect Dis 13: 1840–1846.

2. (2011) Staphylococcus aureus in Healthcare Settings. Centers for Disease Control and Prevention.

3. Lowy FD (1998) Staphylococcus aureus infections. N Engl J Med 339: 520–532.

4. Chambers HF, DeLeo FR (2009) Waves of Resistance: Staphylococcus aureus in the Antibiotic Era. Nat Rev Microbiol 7: 629–641.

5. Mediavilla JR, Chen L, Mathema B, Kreiswirth BN (2012) Global epidemiology of community-associated methicillin resistant Staphylococcus aureus (CA-MRSA). Curr Opin Microbiol 15: 588–595.

6. Nimmo GR (2012) USA300 abroad: global spread of a virulent strain of community-associated methicillin-resistant Staphylococcus aureus. Clin Microbiol Infect 18: 725–734.

7. Edgeworth JD, Yadegarfar G, Pathak S, Batra R, Cockfield JD, et al. (2007) An outbreak in an intensive care unit of a strain of methicillin-resistant Staphylococcus aureus sequence type 239 associated with an increased rate of vascular access device-related bacteremia. Clin Infect Dis 44: 493–501.

8. Tenover FC, McDougal LK, Goering RV, Killgore G, Projan SJ, et al. (2006) Characterization of a strain of community-associated methicillin-resistant Staphylococcus aureus widely disseminated in the United States. J Clin Microbiol 44: 108–118.

9. Holden MT, Feil EJ, Lindsay JA, Peacock SJ, Day NP, et al. (2004) Complete genomes of two clinical Staphylococcus aureus strains: evidence for the rapid evolution of virulence and drug resistance. Proc Natl Acad Sci U S A 101: 9786–9791.

10. Diep BA, Gill SR, Chang RF, Phan TH, Chen JH, et al. (2006) Complete genome sequence of USA300, an epidemic clone of community-acquired meticillin-resistant Staphylococcus aureus. Lancet 367: 731–739.

11. Enright MC, Day NP, Davies CE, Peacock SJ, Spratt BG (2000) Multilocus sequence typing for characterization of methicillin-resistant and methicillin-susceptible clones of Staphylococcus aureus. J Clin Microbiol 38: 1008–1015.

12. Strommenger B, Braulke C, Heuck D, Schmidt C, Pasemann B, et al. (2008) spa Typing of Staphylococcus aureus as a frontline tool in epidemiological typing. J Clin Microbiol 46: 574–581.

13. Aanensen DM, Spratt BG (2005) The multilocus sequence typing network: mlst.net. Nucleic Acids Res 33: W728–733.

14. Zhang K, McClure JA, Elsayed S, Louie T, Conly JM (2005) Novel multiplex PCR assay for characterization and concomitant subtyping of staphylococcal cassette chromosome mec types I to V in methicillin-resistant Staphylococcus aureus. J Clin Microbiol 43: 5026–5033.

15. Ito T, Katayama Y, Asada K, Mori N, Tsutsumimoto K, et al. (2001) Structural comparison of three types of staphylococcal cassette chromosome mec integrated in the chromosome in methicillin-resistant Staphylococcus aureus. Antimicrob Agents Chemother 45: 1323–1336.

16. (2009) Classification of staphylococcal cassette chromosome mec (SCCmec): guidelines for reporting novel SCCmec elements. Antimicrob Agents Chemother 53: 4961–4967.

17. Otto M (2013) Community-associated MRSA: what makes them special? Int J Med Microbiol 303: 324–330.

18. Diep BA, Carleton HA, Chang RF, Sensabaugh GF, Perdreau-Remington F (2006) Roles of 34 virulence genes in the evolution of hospital- and community-associated strains of methicillin-resistant Staphylococcus aureus. J Infect Dis 193: 1495–1503.

19. Li M, Diep BA, Villaruz AE, Braughton KR, Jiang X, et al. (2009) Evolution of virulence in epidemic community-associated methicillin-resistant Staphylococcus aureus. Proc Natl Acad Sci U S A 106: 5883–5888.

20. Fries BC, Varshney AK (2013) Bacterial Toxins—Staphylococcal Enterotoxin B. Microbiol Spectr 1.

21. Cheng J, Wang Y, Cao Y, Yan W, Niu X, et al. (2016) The Distribution of 18 Enterotoxin and Enterotoxin-Like Genes in Staphylococcus aureus Strains from Different Sources in East China. Foodborne Pathog Dis 13: 171–176.

22. Adesiyun AA, Lenz W, Schaal KP (1991) Exfoliative toxin production by Staphylococcus aureus strains isolated from animals and human beings in Nigeria. Microbiologica 14: 357–362.

23. Bukowski M, Wladyka B, Dubin G (2010) Exfoliative Toxins of Staphylococcus aureus. Toxins (Basel) 2: 1148–1165.

24. Ikeda T, Tamate N, Yamaguchi K, Makino S (2005) Mass Outbreak of Food Poisoning Disease Caused by Small Amounts of Staphylococcal Enterotoxins A and H. Appl Environ Microbiol 71: 2793–2795.

25. McCarthy AJ, Lindsay JA (2010) Genetic variation in Staphylococcus aureus surface and immune evasion genes is lineage associated: implications for vaccine design and host-pathogen interactions. BMC Microbiol 10: 173.

26. Omuse G, Kabera B, Revathi G (2014) Low prevalence of methicillin resistant Staphylococcus aureus as determined by an automated identification system in two private hospitals in Nairobi, Kenya: a cross sectional study. BMC Infect Dis 14: 669.

27. Maina EK, Kiiyukia C, Wamae CN, Waiyaki PG, Kariuki S (2013) Characterization of methicillin-resistant Staphylococcus aureus from skin and soft tissue infections in patients in Nairobi, Kenya. Int J Infect Dis 17: e115–119.

28. Aiken AM, Mutuku IM, Sabat AJ, Akkerboom V, Mwangi J, et al. (2014) Carriage of Staphylococcus aureus in Thika Level 5 Hospital, Kenya: a cross-sectional study. Antimicrob Resist Infect Control 3: 22.

29. Omuse G, Shivachi P, Kariuki S, Revathi G (2013) Prevalence of Panton Valentine Leukocidin in Carriage and Infective Strains of Staphylococcus aureus at a Referral Hospital in Kenya. Open Journal of Medical Microbiology Vol.03No.01: 7.

30. Shopsin B, Gomez M, Montgomery SO, Smith DH, Waddington M, et al. (1999) Evaluation of protein A gene polymorphic region DNA sequencing for typing of Staphylococcus aureus strains. J Clin Microbiol 37: 3556–3563.

31. Wattam AR, Abraham D, Dalay O, Disz TL, Driscoll T, et al. (2014) PATRIC, the bacterial bioinformatics database and analysis resource. Nucleic Acids Res 42: D581–591.

32. Zerbino DR, Birney E (2008) Velvet: algorithms for de novo short read assembly using de Bruijn graphs. Genome Res 18: 821–829.

33. Bankevich A, Nurk S, Antipov D, Gurevich AA, Dvorkin M, et al. (2012) SPAdes: A New Genome Assembly Algorithm and Its Applications to Single-Cell Sequencing. J Comput Biol 19: 455–477.

34. Brettin T, Davis JJ, Disz T, Edwards RA, Gerdes S, et al. (2015) RASTtk: a modular and extensible implementation of the RAST algorithm for building custom annotation pipelines and annotating batches of genomes. Sci Rep 5: 8365.

35. Larsen MV, Cosentino S, Rasmussen S, Friis C, Hasman H, et al. (2012) Multilocus sequence typing of total-genome-sequenced bacteria. J Clin Microbiol 50: 1355–1361.

36. Jolley KA, Maiden MC (2010) BIGSdb: Scalable analysis of bacterial genome variation at the population level. BMC Bioinformatics 11: 595.

37. Kuck P, Longo GC (2014) FASconCAT-G: extensive functions for multiple sequence alignment preparations concerning phylogenetic studies. Front Zool 11: 81.

38. Guindon S, Dufayard JF, Lefort V, Anisimova M, Hordijk W, et al. (2010) New algorithms and methods to estimate maximum-likelihood phylogenies: assessing the performance of PhyML 3.0. Syst Biol 59: 307–321.

39. Edgar RC (2004) MUSCLE: multiple sequence alignment with high accuracy and high throughput. Nucleic Acids Res 32: 1792–1797.

40. Stamatakis A (2014) RAxML version 8: a tool for phylogenetic analysis and post-analysis of large phylogenies. Bioinformatics 30: 1312–1313.

41. Wattam AR, Davis JJ, Assaf R, Boisvert S, Brettin T, et al. (2017) Improvements to PATRIC, the all-bacterial Bioinformatics Database and Analysis Resource Center. Nucleic Acids Res 45: D535–D542.

42. Joensen KG, Scheutz F, Lund O, Hasman H, Kaas RS, et al. (2014) Real-time whole-genome sequencing for routine typing, surveillance, and outbreak detection of verotoxigenic Escherichia coli. J Clin Microbiol 52: 1501–1510.

43. Baba T, Bae T, Schneewind O, Takeuchi F, Hiramatsu K (2008) Genome sequence of Staphylococcus aureus strain Newman and comparative analysis of staphylococcal genomes: polymorphism and evolution of two major pathogenicity islands. J Bacteriol 190: 300–310.

44. Johnson AP, Aucken HM, Cavendish S, Ganner M, Wale MC, et al. (2001) Dominance of EMRSA-15 and −16 among MRSA causing nosocomial bacteraemia in the UK: analysis of isolates from the European Antimicrobial Resistance Surveillance System (EARSS). J Antimicrob Chemother 48: 143–144.

45. Hauck CR, Agerer F, Muenzner P, Schmitter T (2006) Cellular adhesion molecules as targets for bacterial infection. Eur J Cell Biol 85: 235–242.

46. Vandenesch F, Lina G, Henry T (2012) Staphylococcus aureus hemolysins, bi-component leukocidins, and cytolytic peptides: a redundant arsenal of membrane-damaging virulence factors? Front Cell Infect Microbiol 2: 12.

47. Omuse G, Van Zyl KN, Hoek K, Abdulgader S, Kariuki S, et al. (2016) Molecular characterization of Staphylococcus aureus isolates from various healthcare institutions in Nairobi, Kenya: a cross sectional study. Ann Clin Microbiol Antimicrob 15: 51.

48. Schaumburg F, Alabi AS, Peters G, Becker K (2014) New epidemiology of Staphylococcus aureus infection in Africa. Clin Microbiol Infect 20: 589–596.

49. Rolo J, Miragaia M, Turlej-Rogacka A, Empel J, Bouchami O, et al. (2012) High genetic diversity among community-associated Staphylococcus aureus in Europe: results from a multicenter study. PLoS One 7: e34768.

50. Aanensen DM, Feil EJ, Holden MT, Dordel J, Yeats CA, et al. (2016) Whole-Genome Sequencing for Routine Pathogen Surveillance in Public Health: a Population Snapshot of Invasive Staphylococcus aureus in Europe. MBio 7.

51. Abdulgader SM, Shittu AO, Nicol MP, Kaba M (2015) Molecular epidemiology of Methicillin-resistant Staphylococcus aureus in Africa: a systematic review. Front Microbiol 6: 348.

52. Driebe EM, Sahl JW, Roe C, Bowers JR, Schupp JM, et al. (2015) Using Whole Genome Analysis to Examine Recombination across Diverse Sequence Types of Staphylococcus aureus. PLoS One 10: e0130955.

53. Strauss L, Stegger M, Akpaka PE, Alabi A, Breurec S, et al. (2017) Origin, evolution, and global transmission of community-acquired Staphylococcus aureus ST8. Proc Natl Acad Sci U S A 114: E10596–E10604.

54. Holden MT, Lindsay JA, Corton C, Quail MA, Cockfield JD, et al. (2010) Genome sequence of a recently emerged, highly transmissible, multi-antibiotic- and antiseptic-resistant variant of methicillin-resistant Staphylococcus aureus, sequence type 239 (TW). J Bacteriol 192: 888–892.

55. Tristan A, Ferry T, Durand G, Dauwalder O, Bes M, et al. (2007) Virulence determinants in community and hospital meticillin-resistant Staphylococcus aureus. J Hosp Infect 65 Suppl 2: 105–109.

56. Holden MT, Hsu LY, Kurt K, Weinert LA, Mather AE, et al. (2013) A genomic portrait of the emergence, evolution, and global spread of a methicillin-resistant Staphylococcus aureus pandemic. Genome Res 23: 653–664.

57. Ruimy R, Maiga A, Armand-Lefevre L, Maiga I, Diallo A, et al. (2008) The carriage population of Staphylococcus aureus from Mali is composed of a combination of pandemic clones and the divergent Panton-Valentine leukocidin-positive genotype ST152. J Bacteriol 190: 3962–3968.

58. Perez-Roth E, Alcoba-Florez J, Lopez-Aguilar C, Gutierrez-Gonzalez I, Rivero-Perez B, et al. (2010) Familial furunculosis associated with community-acquired leukocidin-positive methicillin-susceptible Staphylococcus aureus ST152. J Clin Microbiol 48: 329–332.

59. Monecke S, Slickers P, Ellington MJ, Kearns AM, Ehricht R (2007) High diversity of Panton-Valentine leukocidin-positive, methicillin-susceptible isolates of Staphylococcus aureus and implications for the evolution of community-associated methicillin-resistant S. aureus. Clin Microbiol Infect 13: 1157–1164.

60. Muller-Premru M, Strommenger B, Alikadic N, Witte W, Friedrich AW, et al. (2005) New strains of community-acquired methicillin-resistant Staphylococcus aureus with Panton-Valentine leukocidin causing an outbreak of severe soft tissue infection in a football team. Eur J Clin Microbiol Infect Dis 24: 848–850.

61. Corrigan RM, Rigby D, Handley P, Foster TJ (2007) The role of Staphylococcus aureus surface protein SasG in adherence and biofilm formation. Microbiology 153: 2435–2446.

62. Paharik AE, Horswill AR (2016) The Staphylococcal Biofilm: Adhesins, Regulation, and Host Response. Microbiol Spectr 4.

63. Ramdani-Bouguessa N, Bes M, Meugnier H, Forey F, Reverdy ME, et al. (2006) Detection of methicillin-resistant Staphylococcus aureus strains resistant to multiple antibiotics and carrying the Panton-Valentine leukocidin genes in an Algiers hospital. Antimicrob Agents Chemother 50: 1083–1085.

64. Mbogori C, Muigai A, Kariuki S (2013) Detection and Characterization of Methicillin Resistant Staphylococcus aureus from Toilet and Classroom Door Handles in Selected Secondary Schools in Nairobi County. Open Journal of Medical Microbiology Vol.03No.04: 5.

65. Okon KO, Basset P, Uba A, Lin J, Oyawoye B, et al. (2009) Cooccurrence of predominant Panton-Valentine leukocidin-positive sequence type (ST) 152 and multidrug-resistant ST 241 Staphylococcus aureus clones in Nigerian hospitals. J Clin Microbiol 47: 3000–3003.

66. Varshney AK, Mediavilla JR, Robiou N, Guh A, Wang X, et al. (2009) Diverse enterotoxin gene profiles among clonal complexes of Staphylococcus aureus isolates from the Bronx, New York. Appl Environ Microbiol 75: 6839–6849.

67. Wang CC, Lo WT, Hsu CF, Chu ML (2004) Enterotoxin B is the predominant toxin involved in staphylococcal scarlet fever in Taiwan. Clin Infect Dis 38: 1498–1502.

68. Takahashi N (2003) Neonatal toxic shock syndrome-like exanthematous disease (NTED). Pediatr Int 45: 233–237.

69. Davis JP, Chesney PJ, Wand PJ, LaVenture M (1980) Toxic-shock syndrome: epidemiologic features, recurrence, risk factors, and prevention. N Engl J Med 303: 1429–1435.

70. Shands KN, Schmid GP, Dan BB, Blum D, Guidotti RJ, et al. (1980) Toxic-shock syndrome in menstruating women: association with tampon use and Staphylococcus aureus and clinical features in 52 cases. N Engl J Med 303: 1436–1442.

71. Dinges MM, Orwin PM, Schlievert PM (2000) Exotoxins of Staphylococcus aureus. Clin Microbiol Rev 13: 16–34.

72. Chen TR, Chiou CS, Tsen HY (2004) Use of novel PCR primers specific to the genes of staphylococcal enterotoxin G, H, I for the survey of Staphylococcus aureus strains isolated from food-poisoning cases and food samples in Taiwan. Int J Food Microbiol 92: 189–197.

73. Bing-Shao Liang Y-MH, Yin-Shuang Chen, Hui Dong, Jia-Liang Mai, Yong-Qiang Xie, Hua-Min Zhong, Qiu-Lian Deng, Yan Long, Yi-Yu Yang, Si-Tang Gong and Zhen-Wen Zhou (2017) Antimicrobial resistance and prevalence of CvfB, SEK and SEQ genes among Staphylococcus aureus isolates from paediatric patients with bloodstream infections. Exp Ther Med 14: 5143–5148.

74. Voyich JM, Braughton KR, Sturdevant DE, Whitney AR, Said-Salim B, et al. (2005) Insights into mechanisms used by Staphylococcus aureus to avoid destruction by human neutrophils. J Immunol 175: 3907–3919.

